# Structural characterization of *Myxococcus xanthus* MglC, a component of polarity control system, and its interactions with MglB

**DOI:** 10.1101/2020.08.27.270058

**Authors:** Srajan Kapoor, Akriti Kodesia, Nidhi Kalidas, Ashish, Krishan Gopal Thakur

## Abstract

*Myxococcus xanthus* displays two types of motilities i.e. Social (S) and Adventurous (A). The pole-to-pole reversals of these motility regulator proteins is the key to this process. Here, we determined ~1.85 Å resolution crystal structure of MglC, which revealed that despite sharing <9% sequence identity, both MglB and MglC adopt <u>R</u>egulatory <u>L</u>ight <u>C</u>hain 7 (RLC7) family fold. Interestingly, MglC is structurally unique compared to the other known RLC7 family proteins having ~30°-40° shift in the orientation of functionally important α2 helix. Using isothermal titration calorimetry and gel filtration chromatography, we show that MglC binds MglB in 2:4 stoichiometry with submicromolar range dissociation constant. Using combination of small angle X-ray scattering and molecular docking studies, we show that MglBC complex is formed by MglC homodimer sandwiched between two homodimers of MglB.

**In Brief:** Kapoor *et al*. report the crystal structure of *Myxococcus xanthus* MglC, a <u>R</u>oadblock <u>L</u>ight <u>C</u>hain 7 (RLC7) family protein, involved in polarity reversal. The structure reveals a distinct orientation of α2 helix compared to other RLC7 proteins. They also demonstrate that MglC binds a GTPase activating protein, MglB, with submicromolar range dissociation constant.

**Highlights:**

- MglC adopts RLC7 fold and has distinct structural features.
- MglC interacts MglB to form a stable complex having submicromolar range dissociation constant.
- MglC homodimer is sandwiched between two MglB homodimers to form a 2:4 stoichiometric complex.

## Introduction

*Myxococcus xanthus* is anaerobic, rod shaped gram-negative δ-proteobacteria. It is widely studied for its complex social behaviour, life cycle, and motility (Konovalova et al., 2010; Shi et al., 1993). It exhibits two types of motilities, S “social” motility and A “adventurous” motility (Hodgkin and Kaiser, 1979a, b). In the S-motility, a large group of bacterial cells coordinates to move together. The S-motility uses the type IVa pili filaments that are formed at the leading pole and move the cells forward (Wu and Kaiser, 1995). While, in the A-motility, single cells move at the periphery of bacterial colonies to explore the surrounding environment (Kaiser, 1979). The A-motility is type IVa pilus independent and uses Agl-Glt motility machinery that assembles at the leading pole and provides directionality (Schumacher and Sogaard-Andersen, 2017). The common feature of both types of motility systems is the leading pole assembly of type IVa pili and Agl-Glt complex and their polar inversion by 180° at the opposite poles (Herrou and Mignot, 2020; Islam and Mignot, 2015; Yang et al., 2004). Regulation of pole reversal is essential for modulating motility, which aids adaption and survival in *M. xanthus* (Shapiro et al., 2002). The polar reversals are controlled by “frizzy” signal transduction proteins that act as a “switch control system” (Spormann and Kaiser, 1999; Sun et al., 2000; Zusman, 1982). The Frz system controls the “polarity control system,” i.e. <u>m</u>utual <u>gl</u>iding-motility protein <u>A</u> (MglA), <u>m</u>utual <u>gl</u>iding-motility protein <u>B</u> (MglB), and <u>r</u>equired f<u>o</u>r <u>m</u>otility response regulator complex (RomRX) (Guzzo et al., 2018; Keilberg et al., 2012; Leonardy et al., 2007).

As shown in **Figure 1** before the reversal, MglA, a GTPase, in its GTP bound active form is present at the leading pole (Baranwal et al., 2019; Galicia et al., 2019; Hartzell and Kaiser, 1991). While MglB, a <u>G</u>TPase <u>a</u>ctivating <u>p</u>rotein (GAP), is present at the lagging pole along with RomRX that act as <u>G</u>uanine <u>e</u>xchange <u>f</u>actor (GEF) (Leonardy et al., 2010; Szadkowski et al., 2019; Zhang et al., 2010). The “frizzy” signal transduction proteins part of the Frz system are known to start and regulate the polarity reversal process (Blackhart and Zusman, 1985; Sun et al., 2011). The FrzE phosphorylates its response regulators FrzX and FrzZ that are known to interact and modulate MglB and MglA, respectively (Inclan et al., 2007; Kaimer and Zusman, 2013). The exact sequence of events for the reversal process is not known. However, the MglA proteins first dissociate from the leading pole due to FrzZ signal and travel towards the lagging pole (Herrou and Mignot, 2020). MglA and MglB co-localize at the leading pole for about 30 seconds and in this time the GTPase activity of MglB is not functional due to inactivation by FrzX (Guzzo et al., 2018). Then the MglB proteins detach from lagging pole, move to the opposite pole and RomRX. Load more MglA-GTP molecules to the new leading pole. The RomRX complex then slowly dissociates from the new leading pole and moves to the opposite pole marking the formation of new leading and lagging pole (Herrou and Mignot, 2020). The time taken by RomRX to detach from one pole and get accumulated at the opposite pole in a sufficient amount marks the refractory period as no reversal activity of MglA and MglB can occur at this time (Keilberg and Sogaard-Andersen, 2014).

**Figure 1.**
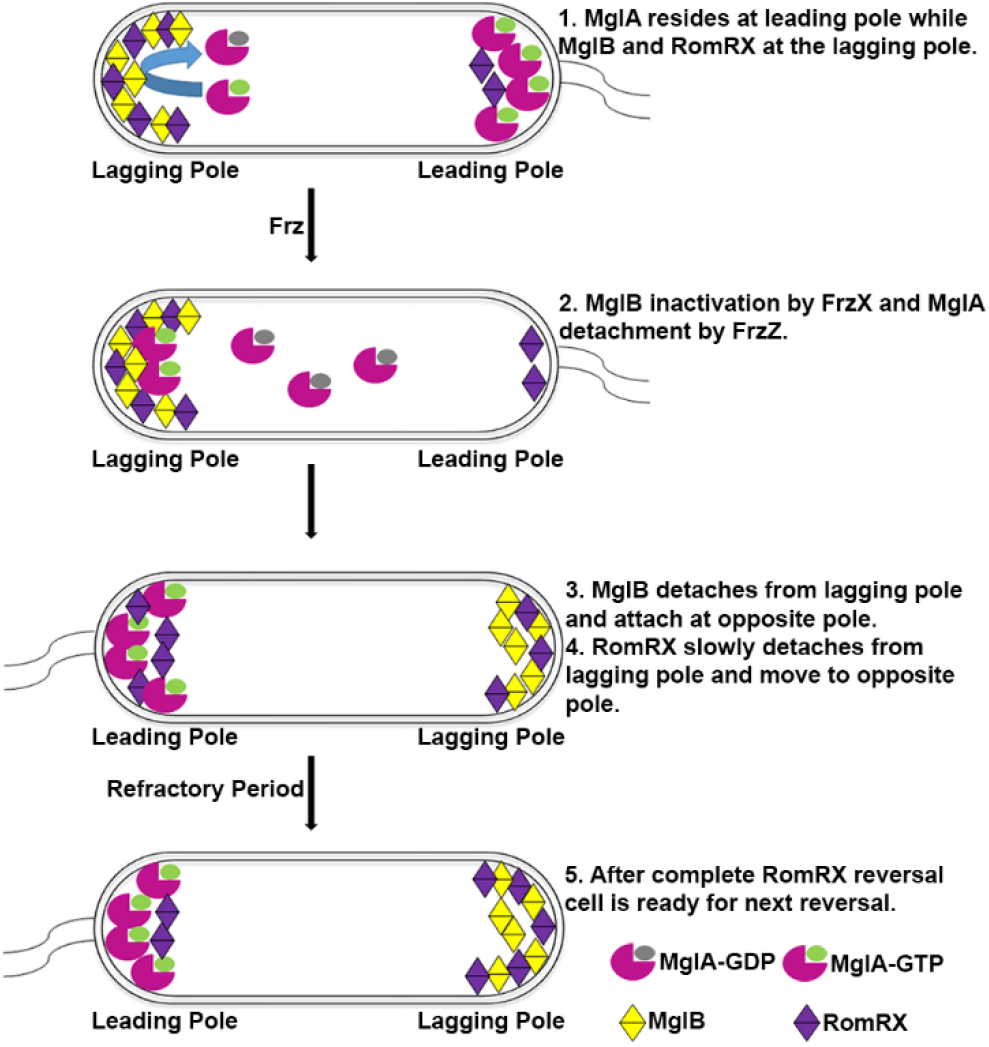
Polar reversal in MglA. Polar reversals of MglA, MglB and RomRX *via* Frz signalling controls the A motility (Gliding motility, Agl-Glt Complex) and S motility (Swarming motility, Type IV pili) of *M. xanthus*.

Currently, the mechanism of the detachment of MglB and RomRX proteins from pole and their attachment to opposite poles is not clear. Recently, a new protein, MglC, has been identified as a member of <u>R</u>egulatory <u>L</u>ight <u>C</u>hain 7 (RLC7) family protein, probably formed by gene duplication and divergence, that over time has lost its ability to bind MglA (McLoon et al., 2016). MglC is required for polarity reversal and interacts with MglB and RomR. MglC is recruited asymmetrically at the lagging poles by RomR as in the absence of RomR the MglC protein is diffused in the cytoplasm (McLoon et al., 2016). MglC is localized in a bipolar manner in the presence of RomR but the presence of MglB mediates localization of MglC mainly at the lagging pole. Using bacterial two hybrid system (BACTH), MglC has been shown to interact with MglB and RomR (McLoon et al., 2016). Upon deletion of MglC, *M. xanthus* cells remain motile but the rate of cellar reversals for motility apparatus reduces significantly suggesting, MglC is probably involved in regulating the rate of cellular reversals for MglB and RomR (McLoon et al., 2016). To understand the role of MglC in regulating pole reversals, it is important to structurally characterize MglC and its interactions with the binding partners.

Here, we determined the crystal structure of MglC at 1.85Å resolution, which revealed structural similarity with MglB despite sharing poor sequence conservation. Comparative structural analysis also revealed distinct structural features compared to other RLC7 family proteins. We further established and characterized MglB and MglC proteinprotein interactions using analytical gel filtration chromatography (GFC) and isothermal titration calorimetry (ITC). Based on data from small angle X-ray scattering (SAXS), gel filtration, ITC, and protein-protein docking studies, we propose a structural model of MglBC protein-protein complex.

## Materials and Methods

### Multiple sequence alignment (MSA)

For multiple sequence alignment based analysis, the protein sequence of MglC (ABF90799.1), was submitted at NCBI BLAST (Altschul et al., 1990) to retrieve sequences sharing sequence similarity. From the BLAST results, ten homologs sharing more than 30% sequence identity were selected for MSA. For comparison with MglB, five MglC and MglB homologs with more than 30% sequence identity were selected for MSA. The MSA was then generated using <u>C</u>onstraint-<u>B</u>ased multiple <u>A</u>lignment <u>T</u>ool (COBALT) (Papadopoulos and Agarwala, 2007). Alignment files and crystal structures MglC and MglB (PDB ID 6HJM) were then used in <u>E</u>asy <u>S</u>equencing in <u>P</u>ostsc<u>ript</u> (ESPript) 3.0 server (Robert and Gouet, 2014) for generating structure-based sequence alignments.

### Cloning, expression and purification of MglC and MglB

The *mglC mglB* and *mglB^ΔCT^* genes were amplified from genomic DNA of *M. xanthus* (DSMZ, Germany, catalogue number 16526) using primers mglC-F (5’-GCTAGTCGCTAGCTCCTTCCGCACGCACCTCGAG-3’), mglC-R (5’-GCTAAAGCTTCTAGAGCTCGGCGCGCACCT-3’), mglB-F (5’-GCTGAAGCTAGCATGGGCACGCAACTGG-3)’, mglB-R (5’-CGTAAAGCTTTTACTCGCTGAAGAGGTTGTCG-3’), mglB^ΔCT^-R (5’-CGTAAAGCTTTTACACCAGGCTCTCGAAGATCTTCGTGAGCTC-3’ synthesized by Sigma-Aldrich, India. The *mglC, mglB* and *mglB^ΔCT^* PCR products were cloned in pET-Duet-A-TEV, (engineered pET-Duet-1, Novagen, vector with a TEV cleavage site) with TEV-cleavable 6x-His tag at N-terminal of the gene cloned between NheI (New England Biolabs Inc.) and HindIII (New England Biolabs Inc.) to yield pET-Duet-A-TEV-mglC, pET-Duet-A-TEV-mglB and pET-Duet-A-TEV-mglB^ΔCT^ clones. Ligated products were transformed in *Escherichia coli* Top10 cells (Novagen, India) and were confirmed by DNA sequencing. Plasmids carrying the desired gene(s) were transformed in *E. coli* Rosetta (DE3) cells (Novagen) and plated on LB agar plate having 100 μg.mL^-1^ ampicillin (Sisco Research Laboratory Pvt. Ltd., India) and 35 μg.mL^-1^ chloramphenicol (Sisco Research Laboratory Pvt. Ltd., India) and were incubated overnight at 37 °C. The colonies obtained on the plates were used for protein purification. A single colony was inoculated in 10 mL LB media and incubated overnight with the constant shaking of 200 rpm at 37 °C for primary culture. 1% of primary culture was inoculated in 750 mL of LB media and induced with 0.3 mM isopropyl β-D-1-thiogalactopyranoside (IPTG) (Gold Biotechnology) at OD_600_ ~ 0.6 and further incubated at 16 °C for 14–16 h with constant shaking at 200 rpm. Cells were harvested by centrifugation at 10000 g for 10 min. Pellet was then re-suspended in 50 mL lysis buffer (20 mM HEPES pH 8.0, 150 mM NaCl). Protease inhibitor cocktail tablets (Roche, Basel, Switzerland) were added to the lysis buffer before sonication. The supernatant was collected after centrifugation at 18,000 g for 30 min. and passed through pre-equilibrated HIS-Select Ni-nitrilotriacetic acid resin (Ni-NTA) (Sigma-Aldrich Co., USA) at 4 °C for binding of 6x-His tagged protein. The protein was then eluted with lysis buffer containing imidazole (Sigma-Aldrich Co., USA) at different concentrations (20 mM, 200 mM and 500 mM) and concentrated using centrifugal ultrafiltration devices (3 kDa cut-off) (Merck India Pvt. Ltd.). This was followed by Gel filtration chromatography using Superdex 200 Increase 10/300 GL column (GE Lifesciences, India) with flow rate of 0.5 mL per min. The desired fractions were pooled and concentrated using centrifugal ultrafiltration devices (Merck India Pvt. Ltd.). The purity and quality of purified protein samples were checked using SDS-PAGE and concentration was measured using bicinchoninic acid (BCA) protein assay (Thermo Scientific, USA).

Purification tag was removed by incubating protein samples with tobacco etch virus (TEV) protease in protease: protein ratio of 1:30 and incubated at 4 °C for 16 hours. The protein digestion was checked on SDS-PAGE, and cleaved protein was further purified by GFC using Superdex 200 Increase 10/300 GL (GE Lifesciences, India) column and concentrated using centrifugal ultrafiltration devices.

### Preparation of selenomethionine-containing protein for phasing experiments

For incorporation of selenomethionine (Sigma-Aldrich Co., USA) into the protein, minimal media was used to grow *E. coli* Rosetta (DE3) cells (Novagen, India) harbouring expression plasmid coding for MglC. The media composed of M9 salts (Na_2_HPO_4_.7H_2_O, 33.97g/L, KH_2_PO_4_, 15 g/L, NaCl, 2.5 g/L, NH_4_Cl, 5 g/L, (Sigma-Aldrich Co., USA), 1M MgSO_4_ (Sigma-Aldrich Co., USA), 1M CaCl_2_ (Sigma-Aldrich Co., USA), 20% glucose (Sigma-Aldrich Co., USA), trace elements and amino acids (threonine 100 mg/L, phenylalanine 100mg/L, lysine 100mg/L, isoleucine 50mg/L, valine 50 mg/L, selenomethionine 60 mg/L) (Sigma-Aldrich Co., USA). All components of minimal media were prepared in Milli Q water and sterilized separately. Glucose, trace elements and amino acids were filter sterilized and all other components were autoclaved. To prepare the media (1 L), 200 mL of M9 salts, 2 mL 1M MgSO_4_, 20 ml glucose, 100 mL 1M CaCl_2_ and 1x trace elements were mixed together and volume was made up to 1 litre using autoclaved water. The primary culture was grown in the similar fashion as mentioned above. The cells from primary culture were harvested in log phase by centrifugation at 4000 rpm for 10 minutes. The pellet was resuspended in minimal media and used as inoculum for secondary culture. The culture was incubated at 37 °C with shaking at 200 rpm. Amino acids were added to the culture at OD_600_ ~ 0.6 and culture was induced by adding 0.3 mM IPTG and incubated at 16 °C for 18-20 hours. Cells were harvested by centrifugation at 9000 g for 10 min. and protein was purified as described in the previous section.

### Crystallization and structure solution

Crystallization trials of MglC were setup with TEV cleaved and un-cleaved protein samples (9 mg/mL, 12 mg/mL and 15 mg/mL). Crystallization trials were performed in three well high throughput crystallization plates (Swissci, Hampton Research, Aliso Viejo, CA, USA) using commercial crystallization screens (Hampton Research, USA and Molecular Dimension, UK). The plates were incubated at 18 °C in Rokimager RI1000 (Formulatrix Inc. USA). Initial hits were observed, after 12 hours of setting up the crystallization trials in various conditions for cleaved, and after two days for uncleaved protein sample. The crystals of uncleaved protein diffracted anisotropically. In contrast, the crystals for TEV cleaved protein diffracted isotropically. We also produced selenomethionine derivatized MglC crystals. The native and selenomethionine SAD data were collected at Elettra synchrotron radiation source, Trieste, Italy. The data were processed using iMOSFLM (Battye et al., 2011) and scaled using AIMLESS in CCP4 software suite (Winn et al., 2011). The crystal structure was determined by Se-single wavelength anomalous diffraction (SAD) method using Phenix.AutoSol module in Phenix software suite (Liebschner et al., 2019). The structure was further improved using several iterative cycles of manual model building in Coot (Emsley and Cowtan, 2004) and refinement using Refmac (Kovalevskiy et al., 2018; Murshudov et al., 2011; Murshudov et al., 1997; Nicholls et al., 2018). The final model had R_Work_/R_Free_ of 15.8/21.5. The detailed data collection statistics, model refinement parameters and validation statistics are provided in **Table 1**.

**Table 1:** Data collection and refinement statistics. Statistics for the highest resolution shell are shown in parentheses

|  | **MglC (PDB ID: 7CT3)** |
| --- | --- |
| **Data collection and processing** |  |
| Beamline | Elettra 11.2C<br>$\lambda=0.97800$ |
| Resolution range (Å) | 27.52 - 1.85 (1.916 - 1.85) |
| Space group | P 6<sub>5</sub> 2 2 |
| a, b, c (Å) | 96.68 96.68 58.28 |
| $\alpha$, $\beta$, $\gamma$ (°) | 90, 90, 120 |
| Total reflections | 367254 (36441) |
| Unique reflections | 14163 (1359) |
| Multiplicity | 25.9 (26.8) |
| Completeness (%) | 99.69 (99.63) |
| Mean I/sigma (I) | 51.73 (2.29) |
| Wilson B-factor | 29.82 |
| R-merge | 0.5634 (2.045) |
| **Refinement** |  |
| R<sub>work</sub>/R<sub>free</sub> | 0.158/0.215 |
| r.m.s.d. (bonds) (Å) | 0.013 |
| r.m.s.d. (angles) (°) | 1.67 |
| Ramachandran favored (%) | 98.31 |
| Ramachandran allowed (%) | 1.69 |
| Number of non-hydrogen atoms | 991 |
| Macromolecules | 921 |
| Ligands | 1 |
| Water | 69 |
| Protein residues | 120 |
| Average B-factor (Å<sup>2</sup>) | 40.06 |
| Macromolecules | 39.09 |
| Ligand | 32.27 |
| Solvent | 53.10 |

### Analytical gel filtration chromatography

MglB, MglC, and MglB/MglC (mixed in different ratios i.e. 1:1, 2:1, and 1:2) were resolved by analytical gel filtration chromatography using Superdex 200 Increase 10/300 (GE Lifesciences, India) column with flow rate of 0.5 mL/min. The absorption was recorded at 280 nm. The data was plotted using Origin 2016 Software suite (OriginLab Corporation, USA).

### Isothermal titration calorimetry (ITC)

ITC experiments were performed at 25°C with 50 μM of MglB in cell titrated with 500 μM of MglC using MicroCal Auto-iTC200 (Malvern MicroCal, LLC, US). Twenty five injections of 0.6 μL (0.1 μL first injection) were given with a spacing of 200 seconds, and the reference power was kept at 10 μCal/sec. The stirring speed of 750 rpm was kept with filter period of 5 seconds. Three control experiments were also setup with the same parameters i.e. titration of buffer alone, titration of MglB with the buffer in a syringe, and buffer titration with MglC in a syringe. Data were analysed using Origin provided with the equipment using one set of sites model (OriginLab Corporation, USA).

### Molecular docking of MglC and MglB

ClusPro 2.0 server (Kozakov et al., 2013; Kozakov et al., 2017; Vajda et al., 2017) was used for docking MglC with MglB. Crystal structures of MglB PDB ID: 6HJM (Galicia et al., 2019) and MglC, determined in this study, were used for the molecular protein-protein docking studies. The top ten docked poses obtained from ClusPro 2.0 server were then further analyzed manually.

### Small Angle X-ray scattering (SAXS)

SAXS experiments were performed using line collimation on SAXSpace Instrument (Anton Paar. Austria) and data were collected at the CMOS Mythen detector (Dectris, Baden, Switzerland). SAXS data was collected on samples of MglB (25, 15 and 10 mg/mL), MglC (35, 15 and 20 mg/mL), and their freshly eluted complex sample from gel filtration i.e. MglBC complex (18, 9 and 13.5 mg/mL) and MglB^ΔCT^C complex (7, 5 and 2 mg/mL), and their matched buffer. For every run ~60 μL of sample or buffer was exposed for 1 hour at 20°C in a thermostated quartz capillary (1 mm). For all datasets, the position of primary beam was corrected using SAXStreat software. Contribution of buffer components were subtracted, data was desmeared using beam profile and finally scattering intensity profile I(q) was saved a function of q, where q is the momentum transfer vector with units in 1/nm. Latter processing were done using SAXSquant software. The I(q) profiles for the proteins and their complexes were then analyzed and processed using ATSAS 3.0.1 and online versions (Franke et al., 2017). Using low q data, Guinier analysis for globular shape profiles was done to estimate Radius of gyration (R_g_), and additionally, using wider q range, distance distribution function was estimated to estimate maximum linear dimension (D_max_) and R_g_ (Putnam, 2016). Considering the presence of flexible or loosely oriented segments, and domains in our protein shapes, we employed chain-ensemble modelling protocol to restore shapes using the SAXS data and deduced parameters using GASBOR program (Svergun et al., 2001). Earlier too, this methodology has been employed to decipher domain-linker shapes (Badmalia et al., 2017; Pandey et al., 2014; Peddada et al., 2013). For GASBOR program, additional inputs were the number of dummy residues (DR) to be used for computing the shape, and were used equal to dimeric state in case of applying P1 symmetry and equal to monomer when considering P2 symmetry. GASBOR jobs were run multiple times to obtain models and all the models were analysed. The MglB (crystal structure), MglC (crystal structure) and MglBC complex (docking models) structures were aligned with the models generated from GASBOR (Svergun et al., 2001) using SUPCOMB (Kozin and Svergun, 2001). For MglBC complex the models obtained by docking were aligned with various GASBOR (Svergun et al., 2001) generated models of MglBC complex and then analysed manually.

### Bioinformatics and structural analysis

PDBePISA (Krissinel and Henrick, 2007) webserver was used for surface area calculations and PDBsum (Laskowski et al., 1997) was used for obtaining protein subunits contact information. MglB and MglC were aligned together using SSM superpose in Coot (Emsley and Cowtan, 2004). PyMOL (The PyMOL Molecular Graphics System, Version 2.0 Schrödinger, LLC) was used to generate molecular graphic figures and to perform r.m.s.d. calculations. Electrostatic surface analysis was performed for using PyMOL Advanced Poisson-Boltzmann Solver (APBS) plugin (Baker et al., 2001). For modeling of conserved residues on MglC structure, ConSurf server was used (Ashkenazy et al., 2016; Ashkenazy et al., 2010; Celniker et al., 2013; Glaser et al., 2003; Landau et al., 2005).

## Results

### Multiple sequence alignment of MglC and MglB reveals distinct sequence features

Based on the multiple sequence alignment (MSA) and predicted secondary structure assignments, MglC has been predicted to be a member of RLC7 family proteins (Levine et al., 2013). Since MglB is also a member of this family sharing ~8% sequence identity and ~17% similarity with MglC, so we performed MSA to identify the conserved regions among these proteins. Though both these proteins share poor sequence identity, as expected we observed only few key highly conserved residues. The representative MSA of MglC and MglB with sequences sharing >30% identity is shown in **Supplementary Figure 1**. All the annotated MglC homologs are shorter compared to MglB due to the absence of the extra N and C-terminal residues **(Supplementary Figure 1a, 1b)**. These extra N-terminal residues of MglB adopt a β-strand conformation while, the C-terminal residues adopt α-helical conformation and linker residue connecting α3-α4. These extra N and C-terminal residues are functionally important in mediating MglA/MglB interactions (Baranwal et al., 2019).

MSA also highlights various other key differences and similarities among MglC and MglB proteins **(Supplementary Figure 1)**. As shown previously by McLoon *et. al*. highly conserved G27 in MglC (structurally equivalent residue G38 in MglB) is crucial for the formation of turn connecting β1-β2 strands (McLoon et al., 2016). We also noticed that G103 in MglC (structurally equivalent residue G112 in MglB) present in α3 is also highly conserved. The residue at 106 position in MglC (115 in MglB) is also occupied by positively charged amino acid in both MglC and MglB. In MglC, we also observed another conserved G67, which is absent in MglB. In MglC, the residue number 62 is occupied by negatively charged residues while in MglB this equivalent position 79 is occupied by positively charged residue. It has been previously shown that F25, D26 and I28 (FDI sequence motif) of MglC might be involved in the MglBC interactions (McLoon et al., 2016). Our MSA analysis suggests that highly conserved I28 residue of MglC, is absent in MglB. The MglC is characterized by the presence of negatively charged [D/E]26 but is not conserved in MglB. However, as also observed by McLoon *et. al*. there is the presence of conserved D36 in MglB near this position (McLoon et al., 2016).

### Crystal structure of MglC

We successfully crystallized and determined the crystal structure of MglC. MglC crystallized in P6522 space group (a=96.69Å, b=96.69Å, c=58.28Å, α=90°, β=90°, γ=120°) with one molecule in an asymmetric unit. There was no good structural template to use as a model for solving structure using molecular replacement. The closest homologue available at RCSB PDB shared ~8% sequence identity. So, we solved the crystal structure using Se-SAD experimental phasing technique at 1.85Å resolution. The final refined model contains all 120 residues of MglC. The crystal structure revealed the typical RLC7 fold (α1β1β2α2β3α3) **(Figure 2A)**. MglC monomer contains five antiparallel β-strands sandwiched between three α-helices. MglC also contains a 310 helix just after the α2 helix. The two β hairpins connect β1-β2 and β4-β5 strands. Other secondary structural elements include five β-bulges and eight β turns. We also observed an electron density for a metal ion in the crystal structure having square pyramidal geometry. We placed Mg2+ ion which is coordinated by three water molecules and main chain oxygen atoms of T66, and V69.

**Figure 2.**
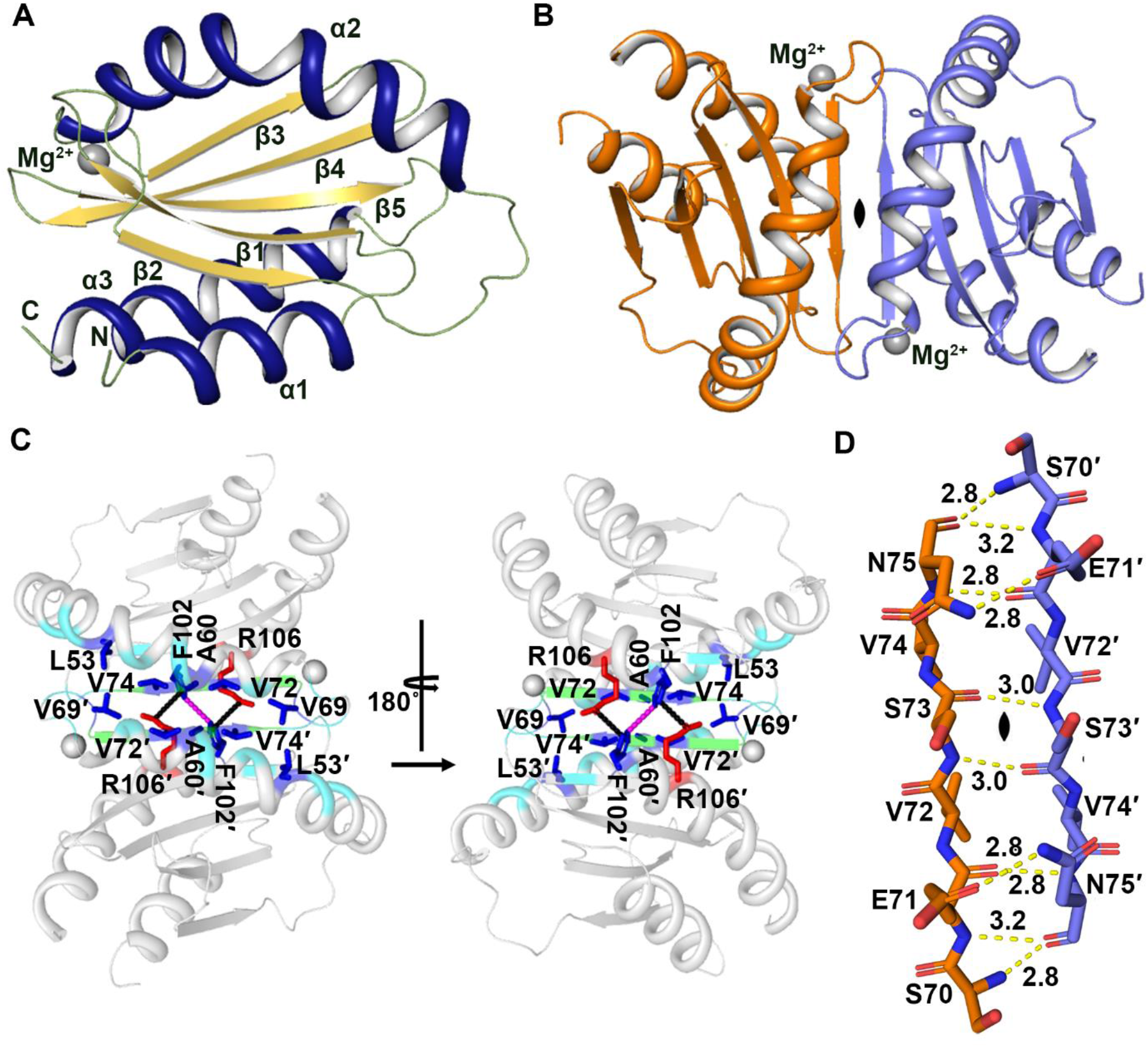
Structural analysis of MglC. **(A)** Cartoon representation of MglC monomer (asymmetric unit) colored according to the secondary structural elements (helices – deep blue, β-sheets – yellow orange, loops and turns – smudge). **(B)** The biological assembly of MglC generated by applying two-fold crystallographic symmetry. The protomers are shown in orange and slate colours. **(C)** The detailed view of the dimeric interface. Residues involved in hydrophobic interactions are shown in blue sticks, R106 and F102 involved in cation-π interaction (black dotted line) are highlighted in red and blue sticks, respectively. The F102 from both protomers are involved in aromatic-aromatic interaction shown in magenta dash line. The β sheet formed at the interface stabilized by main chain hydrogen bonds is shown in green colour. The remaining residues at the interface are shown in cyan **(D)** The hydrogen bonds formed by main chain and side chain interactions at the β sheet extension are shown in yellow lines.

The structural analysis further suggests that, like other members of RLC7 family, MglC also forms homodimer and two protomers are related by crystallographic two-fold symmetry **(Figure 2B, Supplementary Figure 2A)**. PDBePISA (Krissinel and Henrick, 2007) analysis suggests that MglC monomer has total surface area of 6277 Å^2^ and the dimer interface buries 792.1 Å^2^ area (~12.61%). The total surface area of MglC dimer is 10970 Å^2^ with the 1580 Å^2^ (~14.4%) being buried. The ΔG^int^ predicted using PDBePISA (Krissinel and Henrick, 2007) for the dimer association is −9.1 kcal/mol and ΔG^diss^ is 3.9 kcal/mol suggesting that MglC may form a stable dimer. MglC dimerization, is mediated by β sheet extension i.e. β3 from each monomer comes together to form an antiparallel sheet consisting of ten β strands (five strands from each monomer) sandwiched by four helices on one side and two helices on the other side. The dimer is stabilized by six main chain hydrogen bonds and four hydrogen bonds involving side chains between the interacting beta strands at binding interface. Besides these, several non-bonded interactions are also observed between the interacting β strands and α2 helices at the binding interface **(Figure 2C, Supplementary Figure 2B)**. The residues involved in the formation of main chain H-bonds at the dimeric interface i.e. S70, E71, S73, V74, and N75 are highly conserved in MglC. Residue N75 and E71 also form hydrogen bonds with E71 and N75 of the other monomer **(Figure 2D, Supplementary Figure 2B)**. The α2 helices form hydrophobic contacts between chain A and chain B *via* the residues L53-V69’, A60-A60’, V69-L53’, V69-V74’, V72-V74’, V74-V69’, V74-V72’ and F102-F102’. The F102-F1012 is also involved in aromatic-aromatic interactions. Besides this F102 and R106 form a cation-Π interaction **(Figure 2C, Supplementary Figure 2B)**. To further study the oligomeric status of MglC in solution we performed analytical gel fiteration experiments. MglC eluted predominantly at ~17 mL corresponding to molecular weight of ~35 kDa which is close to the expected size of the dimer (~32 kDa). This further supports the crystallographic observations hence confirming that MglC predominantly exist as a dimer in solution **(Supplementary Figure 2A)**.

To obtain information about the probable binding sites of MglC with MglB and RomR, we preformed cleft analysis of MglC using PDBsum (Laskowski et al., 1997). Our analysis revealed the presence of 3 prominent clefts in MglC **(Figure 3A)**. Cleft 1 is largest among all and is formed at the interface of the dimer with a volume of ~3946 Å^3^. This cleft 1 includes highly conserved positively charged residues imparting a net positive charge to the region as revealed by the ConSurf analysis and electrostatic potential map, respectively **(Figure 3B, 3C)** (Ashkenazy et al., 2016; Ashkenazy et al., 2010; Baker et al., 2001; Celniker et al., 2013; Glaser et al., 2003; Landau et al., 2005). The cleft 2 and 2’ (prime is used for identical cleft on another protomer in dimer) includes ‘FDI’ region shown to be involved in binding MglB and has a volume of ~580 Å^3^. The clefts 3 and 3’ have ~600 Å^3^ volume and are lined by variable residues. So, these clefts may be the probable sites for mediating interactions with the binding partners. ConSurf (Ashkenazy et al., 2016; Ashkenazy et al., 2010; Celniker et al., 2013; Glaser et al., 2003; Landau et al., 2005) and MSA analysis revealed high conservation in the turn region (F26 to I29) connecting β1 −β2 strands and these residues (F26, D27 and I29) have been previously shown to be involved in binding MglB **(Figure 3B, 3C)**. The other highly conserved regions are highlighted in the **Figure 3B**.

**Figure 3.**
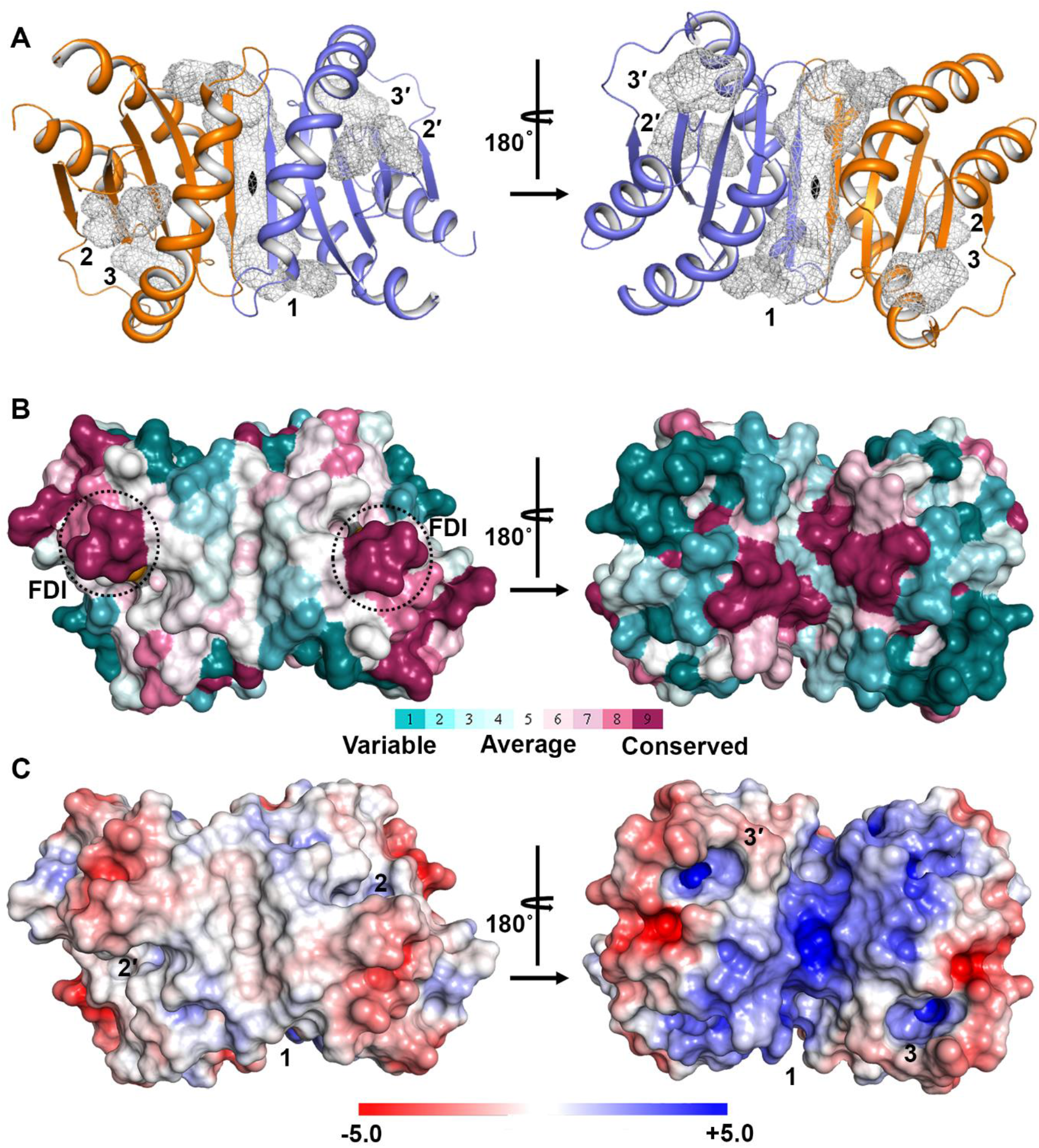
Surface analysis of MglC. (A) Cleft analysis of MglC shows the presence of 3 clefts in MglC. Cleft 1 is largest and is formed at the interface of dimer with the volume of ~3946 Å^3^. This cleft mainly contains highly conserved positively charged residues. **(B)** Consurf analysis (Ashkenazy et al., 2016; Ashkenazy et al., 2010; Celniker et al., 2013; Glaser et al., 2003; Landau et al., 2005) showing sequence conservation mapped on to the MglC crystal structure. The highly conserved region marked by black dotted circle includes ‘FDI’ sequence motif previously shown to be involved in interacting with MglB. **(C)** Electrostatic surface potential of MglC showing distribution of negatively (red), and positively (blue) charged regions (Baker et al., 2001).

### MglC has distinct structural features among RLC7 family proteins

We searched for proteins sharing structural similarity with MglC using PDBeFold (Krissinel and Henrick, 2004). The top hit included late endosomal/lysosomal <u>a</u>daptor and <u>M</u>APK and M<u>TOR</u> activator 2 (LAMTOR2) protein having 1.87 Å r.m.s.d. over 94 residues sharing 10% sequence identity. Interestingly, MglB shares ~8% sequence identity and is among the top hits having 2.0 Å r.m.s.d. over 95 residues. The top hits obtained from the PDBeFold (Krissinel and Henrick, 2004) and structural comparisons are provided in **Supplementary Table 1, 2**. MglB α2 is functionally important and has been shown to interact with switch 1 and switch 2 regions of MglA (Baranwal et al., 2019; Galicia et al., 2019). Comparative structural analysis revealed that α2 in MglC is drastically shifted as compared to MglB and other RLC7 fold proteins **(Figure 4, Supplementary Figure 3)**. For example, α2 helix of MglC is tilted by ~40° and ~33° having maximal displacement of 17.7 Å and 13.8 Å compared to α2 of MglB (6HJM, *M. xanthus)* and LAMTOR2 (5Y3A, *Homo sapiens)* (Galicia et al., 2019; Zhang et al., 2017), respectively **(Figure 4)**. The relative shift in α2 of MglC compared to α2 of other RLC7 family proteins is given in **Supplementary Table 1**. In addition, there is a presence of 310 helix connecting α2 and β2 and highly conserved G67 that is present only in MglC. Upon comparison of MglC dimer with the dimers of other RLC7 family members we also observed that α1 and α3 helices are shifted slightly inward in MglC. For example, compared to *M. xanthus* MglB α1, MglC α1 is shifted by an angle of ~7.2° and displacement of 3.64 Å while, compared to *H. sapiens* LAMTOR2 α1 it is shifted by an angle of ~14.6° and displacement of 4.86 Å. The α3 helix of MglC is also shifted by an angle of ~15.9° and ~10.7° and displacement of 2.73 Å and 3.44 Å compared to *M. xanthus* MglB and *H. sapiens* LAMTOR2 **(Supplementary Figure 3)**. This comparative structural analysis reveals similarities and distinct structural features in MglC compared to other members of RLC7 family proteins. The comparative r.m.s.d. analysis of MglC homodimer with dimers of RLC7 family proteins is shown in **Supplementary Figure 3** and **Supplementary Table 2**.

**Figure 4.**
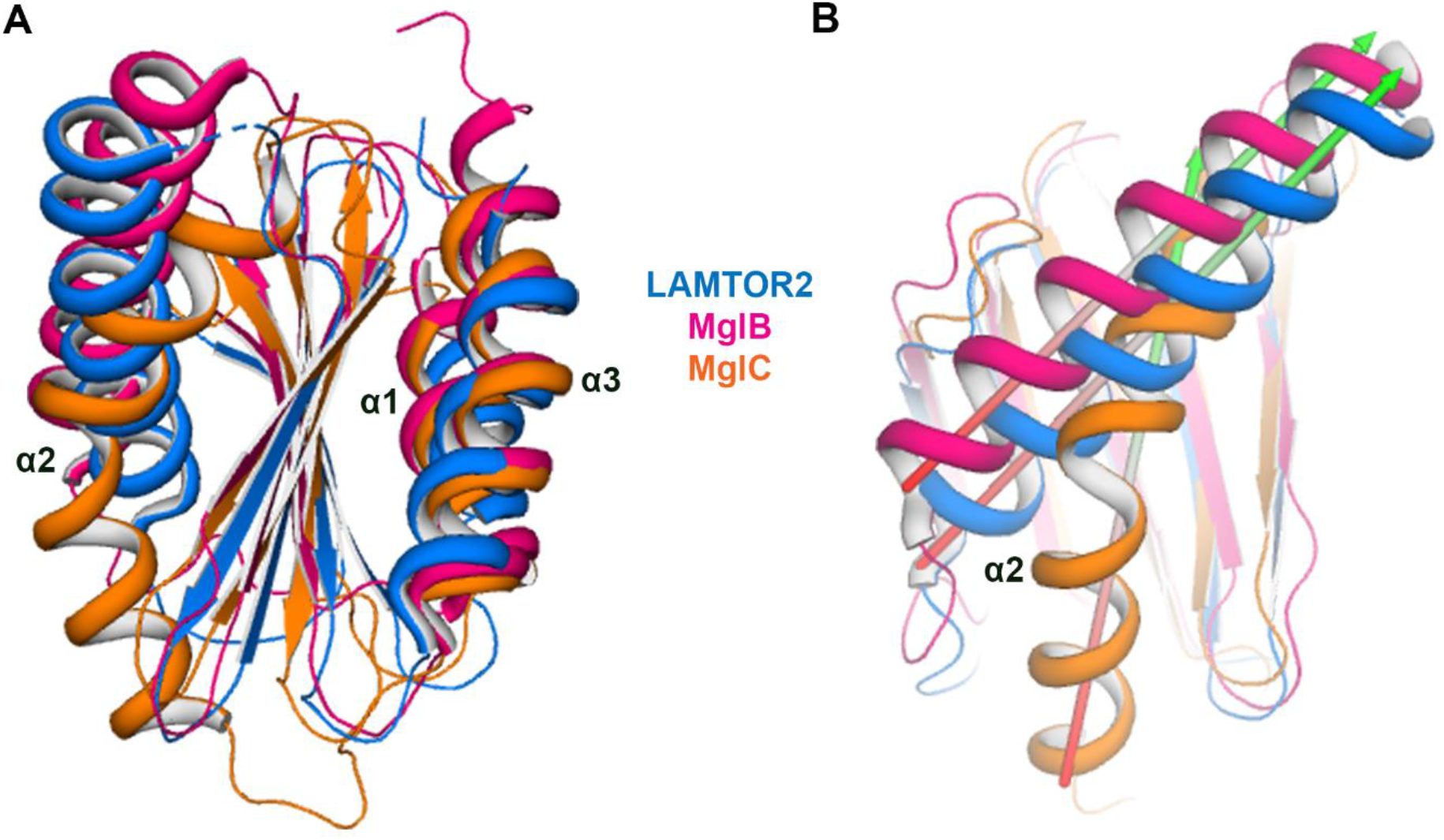
Comparative structural analysis of MglC with other RLC7 family proteins. **(A)** Structural superposition of MglC in orange with other proteins of RLC7 family LAMTOR2 (Zhang et al., 2017) (PDB ID: 5Y3A, Blue), and MglB from *M. xanthus* (Galicia et al., 2019), (PDB ID 6H5B, Pink). **(B)** Structural comparison of MglC showing difference in relative orientation of α2 compared to other RLC7 family proteins.

### MglC interacts with MglB with submicromolar range dissociation constant

Using BACTH assay, McLoon *et. al*. have proposed that MglC might interact with MglB via the ‘FDI’ interface (McLoon et al., 2016). To check physical interactions between MglB and MglC we mixed both the purified proteins in varying ratios and performed analytical gel filtration experiments.. MglB and MglC eluted predominantly at ~15 mL and ~17 mL corresponding to the observed molecular weights of ~42 kDa and ~35 kDa (MglC, Mol. Wt. 32 kDa; MglB, Mol. Wt. 40 kDa) suggesting both proteins exist in homodimeric oligomeric state in solution **(Figure 5A)**. When we mixed both the proteins in the equimolar ratio, we obtained two peaks corresponding to MglBC complex at ~13 mL and excess MglC alone at 17 mL confirming interactions of MglC with MglB **(Figure 5A)**. When the MglB:MglC were mixed in 2:1 ratio, we observed only a small MglC peak and while the majority of the sample eluted as a MglBC complex. However, when MglB: MglC were mixed in 1:2 ratio there was no increase in the intensity of MglBC complex peak while excess MglC peak was observed at 17 mL **(Figure 5A)**. This suggested that MglC binds MglB in 2:4 stoichiometry (as both proteins exist in homodimeric state in solution) to form MglBC complex. MglBC complex was stable during the gel filtration run, suggesting MglC binds MglB with high affinity. Therefore, we performed ITC experiments to determine binding affinity and stoichiometry of MglB-MglC interactions. We used one site model for data fitting. ITC data revealed Kd of ~625 nM and stoichiometry (N) of ~0.5, hence confirming that two homodimeric MglB molecules bind one homodimeric MglC **(Figure 5B)**. As the enthalpy (ΔH) is positive 6944 cal/mol i.e. endothermic process along with positive entropy (ΔS) 51.7 cal/mol/K it suggests that the reaction is entropy-driven.

**Figure 5.**
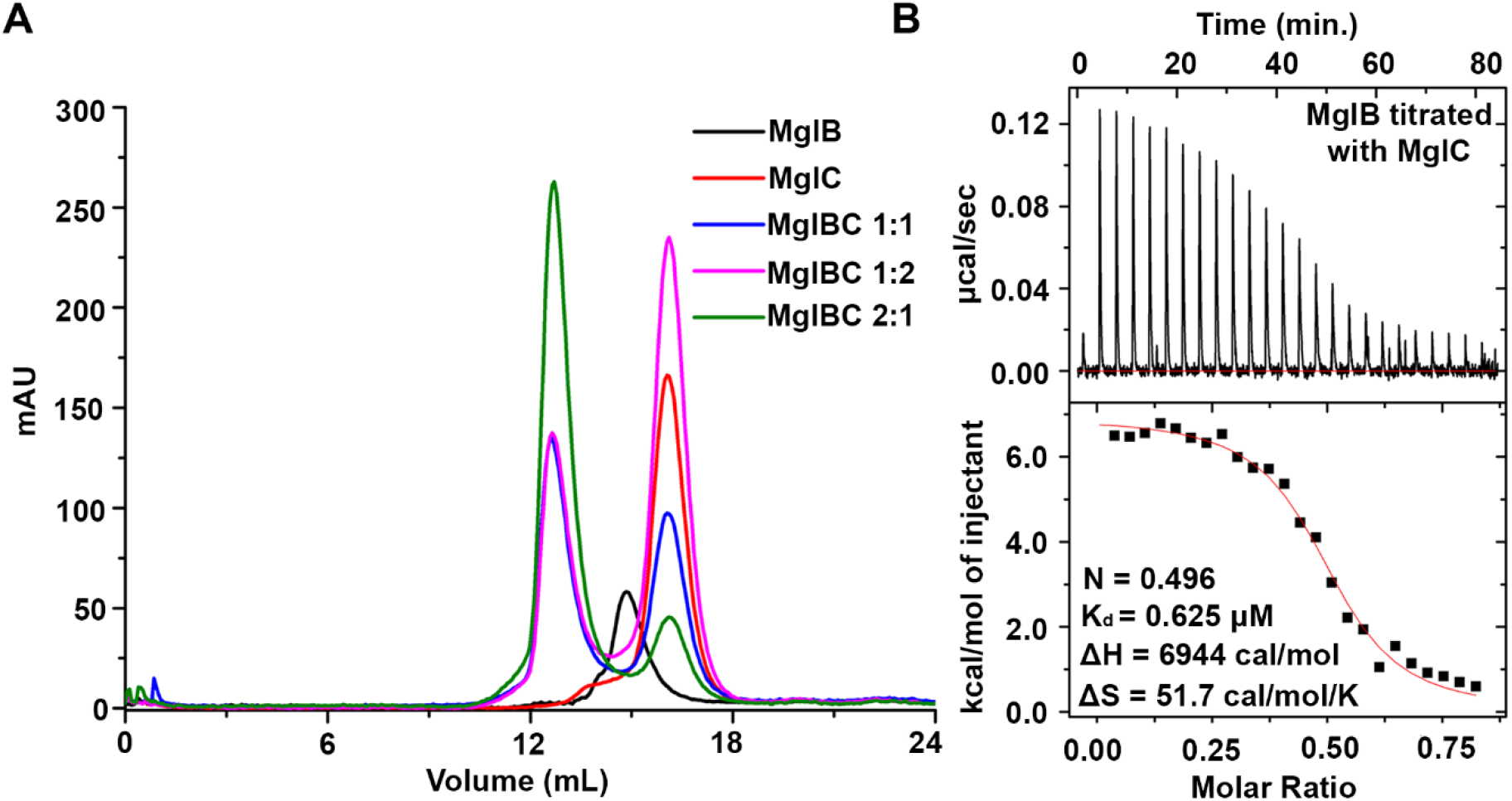
Interaction of MglC with MglB. **(A)** Analytical gel filtration chromatography profile showing the interaction of MglB with MglC. **(B)** Isothermal titration calorimetry profile of MglB-MglC interactions.

### The C-terminal region of MglB is not involved in binding MglC

Crystal structure of MglAB complex revealed involvement of the C-terminal region (residue 147-157 of MglB protomer) of MglB in binding MglA. This was further confirmed by deletion studies (Baranwal et al., 2019). In MglAB complex, the C-terminal region of only one protomer is involved in binding MglA (Baranwal et al., 2019). To investigate the role of this C-terminal region of MglB in binding MglC, we created a deletion variant of MglB (MglB^ΔCT^). We performed gel filtration-based protein-protein interaction studies using this deletion variant. Our data suggests that the MglB deletion construct retains the ability to bind MglC **(Supplementary Figure 4)**. This data suggests the MglB may adopt a distinct mode of binding MglC to form MglBC complex compared to MglAB complex.

### MglC is sandwiched between two MglB molecules

The studies presented above demonstrated that MglBC forms a stable complex in solution. We performed extensive crystallization experiments to determine the structure of MglBC complex. We successfully crystallized the complex however, we did not succeed in improving the diffraction quality of the crystals. So, we next performed protein-protein docking using the ClusPro 2.0 server. Several models were generated by this server. The top ten models were shortlisted for manual analysis. McLoon *et.al*. have previously shown that MglB binds the ‘FDI’ surface of MglC (McLoon et al., 2016). Interestingly, in all the docked structures from ClusPro 2.0 server (Kozakov et al., 2013; Kozakov et al., 2017; Vajda et al., 2017), we observed that MglB was indeed docked at the FDI interface on MglC **(Supplementary Figure 5)**. ‘FDI’ interface is a part of cleft 2 as described in the previous section. Also, none of the models obtained from the docking servers showed involvement of the C-terminal region of MglB in binding MglC which correlates well with our experimental findings. Based on the experimentally determined binding stoichiometry and docking results we generated a model of MglBC complex where MglC homodimer is sandwiched between the two MglB homodimers **(Supplementary Figure 6).** In this proposed model, the MglB homodimer interacts with one chain of MglC and the complex is related by two-fold symmetry.

### SAXS based low resolution in solution structure of MglBC complex

To further confirm oligomeric status, binding stoichiometry and predicted MglBC complex model we performed SAXS experiments. We collected SAXS data on three different concentrations of unliganded MglB, and MglC, and their complex, MglBC as described in the methods section. All data collected for the samples were free from inter-particle effects or aggregation. Guinier analysis considering globular scattering profile estimated R_g_ values of 3.35 nm, 1.99 nm and 3.69 nm for MglB, MglC and MglBC, respectively **(Figure 6A)**. Dimensional Kratky analysis of all the proteins suggested that all the proteins were folded and first maxima was at 1.7 for unliganded MglC and MglBC, and for MglB, the maxima was >1.7 implying partly disordered portion in its shape **(Figure 6B)**. D_max_ obtained for MglB, MglC, and MglBC were 11.5 nm, 6.8 nm, and 14.5 nm, respectively **(Figure 6C)**. The molecular weight was calculated by dividing the Porod volume (~89,089 Å^3^, ~47,239 Å^3^ and ~207,733 Å^3^ for MglB, MglC and MglBC, respectively) by 1.7. This calculated molecular weight to be ~52 kDa, ~28 kDa and ~122 kDa of MglB, MglC and MglBC, respectively which were found to be in close agreement with the theoretical molecular weights for homodimeric MglB (~40 kDa), homodmeric MglC (~32 kDa) and 2:4 stoichiometric MglBC complex (~112 kDa). The dummy-atom models were then build using GASBOR program (Svergun et al., 2001). The fitting of one of the GASBOR generated model with intensity profile is shown in **Figure 6D.** These models were then aligned to the crystal structures of MglC and MglB and molecular docking generated models via SUPCOMB program which aligns inertial axes of the models (Kozin and Svergun, 2001) **(Figure 7)**. Residue resolution structure of MglC aligned well with the shape profile solved using the SAXS data based constraints **(Figure 7A)**. However, models generated for MglB and MglBC complex revealed extra regions which were not resolved in the crystal structure of MglB, probably due to inherent flexibility or accessibility to different local conformations relative to structure domain **(Figure 7B, 7C)**. To further investigate the role of the C-terminal region in proteinprotein interactions in MglBC complex we performed SAXS experiments using MglB^ΔCT^ as well. The SAXS data suggests that MglB^ΔCT^C forms a complex with D_max_ of 10.4 nm **(Figure 7D and Supplementary Figure 7)** further supporting data obtained using gel filtration chromatography. So, the SAXS experiments confirmed the oligomeric states of the MglB, MglC and MglBC protein-protein complex. Also, these experiments further strengthens a molecular docking based model proposed for MglBC complex, where probably MglC homodimer is sandwiched between the two MglB homodimers **(Supplementary Figure 6)**.

**Figure 6.**
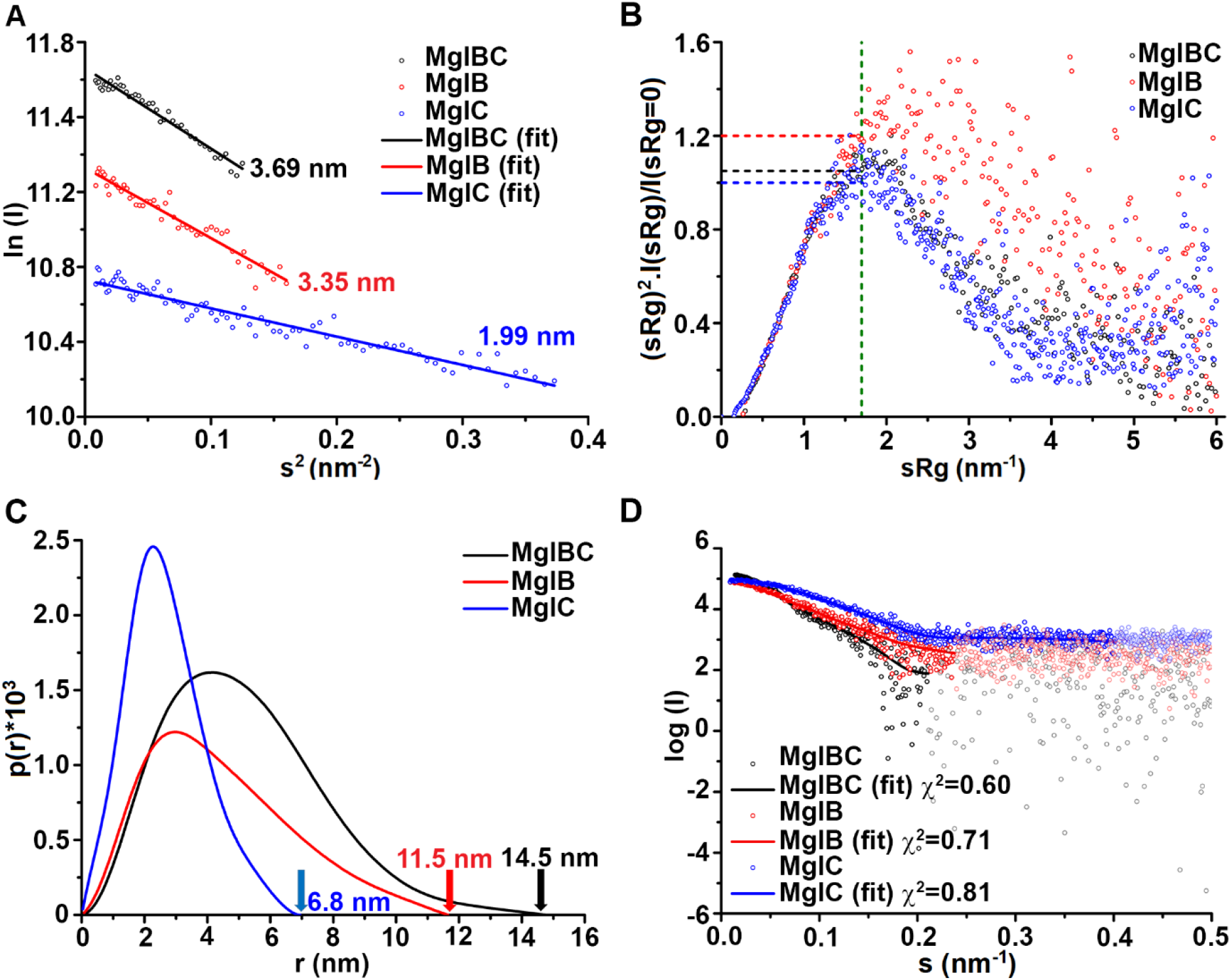
SAXS analysis of MglC (Blue), MglB (Red) and MglBC complex (Black). **(A)** Guinier analysis of MglC (Rg = 1.99 nm), MglB (Rg = 3.35 nm) and MglBC (Rg = 3.69 nm) complex reveals linear fit with no signs of inter-particle effects. **(B)** Dimensionless Kratky plots of MglC and MglBC complex revealed globular nature of these proteins. Kratky plot revealed that the MglB was folded and had some flexible regions. **(C)** Normalized pair distribution function P(r) analysis revealed D_max_ of ~6.8 nm for MglC, 11.5 nm for MglB and ~14.5 nm for MglBC complex. **(D)** Dummy atom model for MglC (χ^2^ = 0.81), MglB (χ^2^ = 0.71) and MglBC (χ^2^ = 0.60) complex were prepared using GASBOR (Svergun et al., 2001). The intensity profile of all three proteins are shown in spheres and the lines represent the fitting of dummy atom models generated by GASBOR (Svergun et al., 2001).

**Figure 7.**
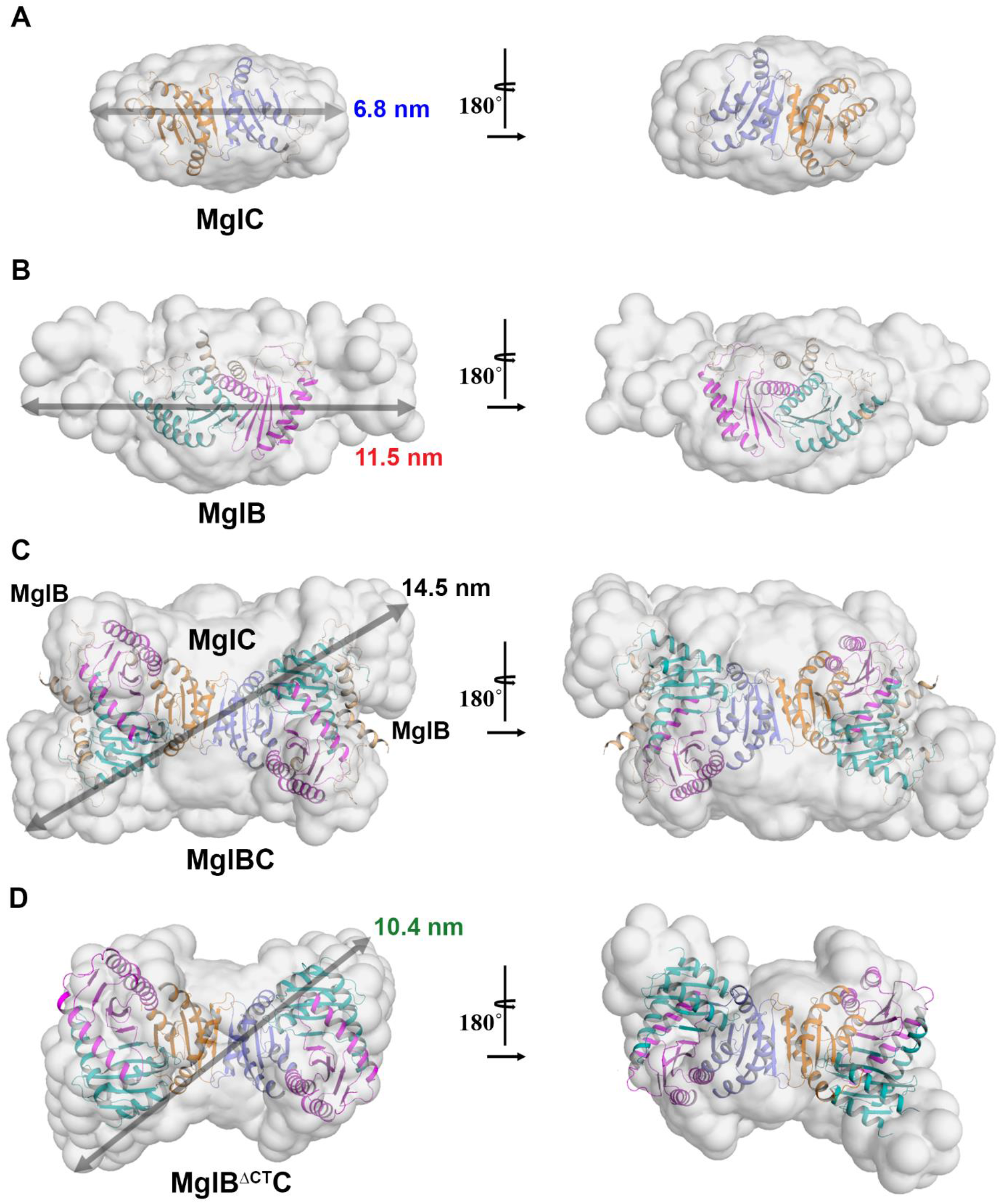
Low resolution solution structure of MglC, MglB and MglBC complex fitted in the SASX envelope using high resolution crystals structures and modelled regions not resolved in crystal structures. **(A)** MglC homodimer superposes well in the dummy model **(B)** The MglB homodimer core fits well. The extra regions in the SAXS envelope are probably due to the flexible N- and C-terminal regions in MglB. **(C)** Low resolution structure of MglBC complex obtained by molecular docking of MglC with MglB reveals the presence of MglC dimer sandwiched between two molecules of MglB dimer. The high-resolution models were fitted in the low-resolution models using SUPCOMB (Kozin and Svergun, 2001). The unaccounted regions in the dummy atom model suggest the presence of flexible regions not resolved well in the crystal structures.

## Discussion

Cellular polarity reversal is an important phenomenon required for various essential functions for existence like motility, development, biofilm formation and many more (Herrou and Mignot, 2020; Mauriello and Zusman, 2007). MglC is newly discovered and least studied member involved in polarity reversal. There is only one publication describing the discovery and role of MglC in polar reversal. In this study, we determined the crystal structure of a recently identified member of polarity reversal complex, MglC. Crystal structure MglC revealed structural similarity with MglB hence confirming both are members of the RLC7 protein family. Despite sharing similar fold architecture, there are distinct structural features in MglC, including deletion of N-and C-terminal regions, differences in the overall arrangement of the secondary structural elements which might dictate preferences for specific binding partners. We also show that MglC binds MglB with 2:4 stoichiometry respectively, to form a stable complex with submicromolar range dissociation constant.

Cleft analysis of MglC suggested presence of three distinct clefts i.e. 1, 2 (and 2’) and 3 (and 3’) in each protomer. Molecular protein-protein docking results obtained from ClusPro 2.0 (Kozakov et al., 2013; Kozakov et al., 2017; Vajda et al., 2017) suggested involvement of cleft 2 and 2’ in binding MglB. We used SAXS based analysis to further obtain low resolution insights to propose a model for MglBC complex. MglC cleft 2 contains highly conserved ‘FDI’ sequence motif which has been shown to be involved in binding MglB (McLoon et al., 2016).

MglB reportedly binds to several proteins involved in polar reversal including MglA, RomRX, SofG and MglC (Baranwal et al., 2019; Galicia et al., 2019; Kanade et al., 2020; McLoon et al., 2016; Szadkowski et al., 2019). MglAB complex has been structurally characterized (Baranwal et al., 2019; Galicia et al., 2019). Based on pull-down experiments of 6x-His tagged MglB with *M. xanthus* cell lysate it has been shown that MglB and RomR could potentially interact (Keilberg et al., 2012; Zhang et al., 2012). MglC also reportedly interacts with RomR, but probably at the binding site different from MglB (McLoon et al., 2016). All these studies and the data presented here suggest that both MglB and MglC have multiple binding partners, and partner switching may be required for polar reversal. To our knowledge, the binding affinities for these reported binding partners with MglB or MglC has not yet been determined. Hence we could not compare affinities with other known complexes involved in polarity reversal. In our binding studies we observed moderate submicromolar dissociation constant for MglBC complex. We speculate that this moderate affinity may facilitate partner switching at cellular concentrations. MglA (small GTPase), MglB (GAP) and RomR (GEF) are known to be modulated by Frz mediated phosphorylation (Guzzo et al., 2018; Keilberg et al., 2012; Szadkowski et al., 2019). It could be possible that Frz signalling also play significant role in modulating the binding partners of MglC or could modulate the binding affinity of MglC for MglB and RomR. This needs a thorough investigation in the future studies.

To summarize, we report the first structural description of MglC involved in polarity reversal from *M. xanthus*. We have established and characterized MglBC complex in detail. Our data suggests that MglC exists in homodimeric oligomeric state in solution and interacts with two molecules of MglB homodimers with submicromolar range dissociation constant. In future, it will be interesting to know how cellular reversals are regulated by protein-protein interactions involving multiple binding partners. Detailed structural and functional studies will be required to understand the role of protein-protein interactions in mediating polar reversal.

## Acknowledgments

Authors thank Mr. Surinder Singh at CSIR-IMTECH Chandigarh for supporting Laboratory experiments. KGT would like to acknowledge members of Structural Biology Laboratory for useful suggestions and discussions.

