## Supplementary data for "Structural characterization of *Myxococcus xanthus* MglC, a component of polarity control system, and its interactions with MglB"

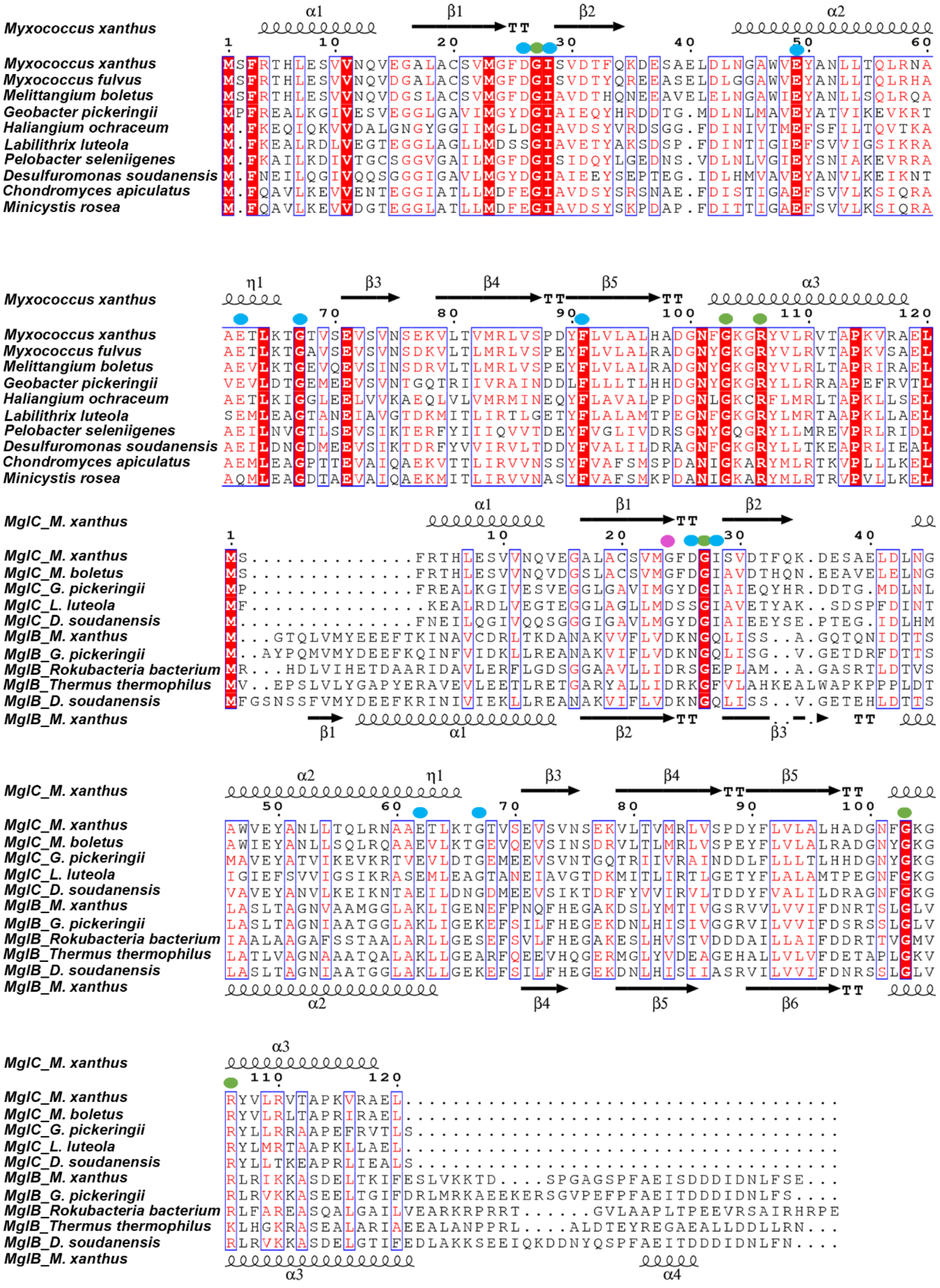


**Supplementary Figure 1: Multiple Sequence alignment (A)** Multiple Sequence Alignment of MglC homologs with more than 30% sequence identity **(B)** Multiple sequence alignment of MglC (more than 30% sequence identity) and MglB (more than 30% sequence identity) highlighting the conserved residues in MglB and MglC sequence by Green dots. The highly conserved residues in MglC that are mutated in MglB are marked by blue dots. The pink dot mark the residue conserved in MglB but mutated in MglC. (Altschul et al., 1990; Papadopoulos and Agarwala, 2007; Robert and Gouet, 2014).

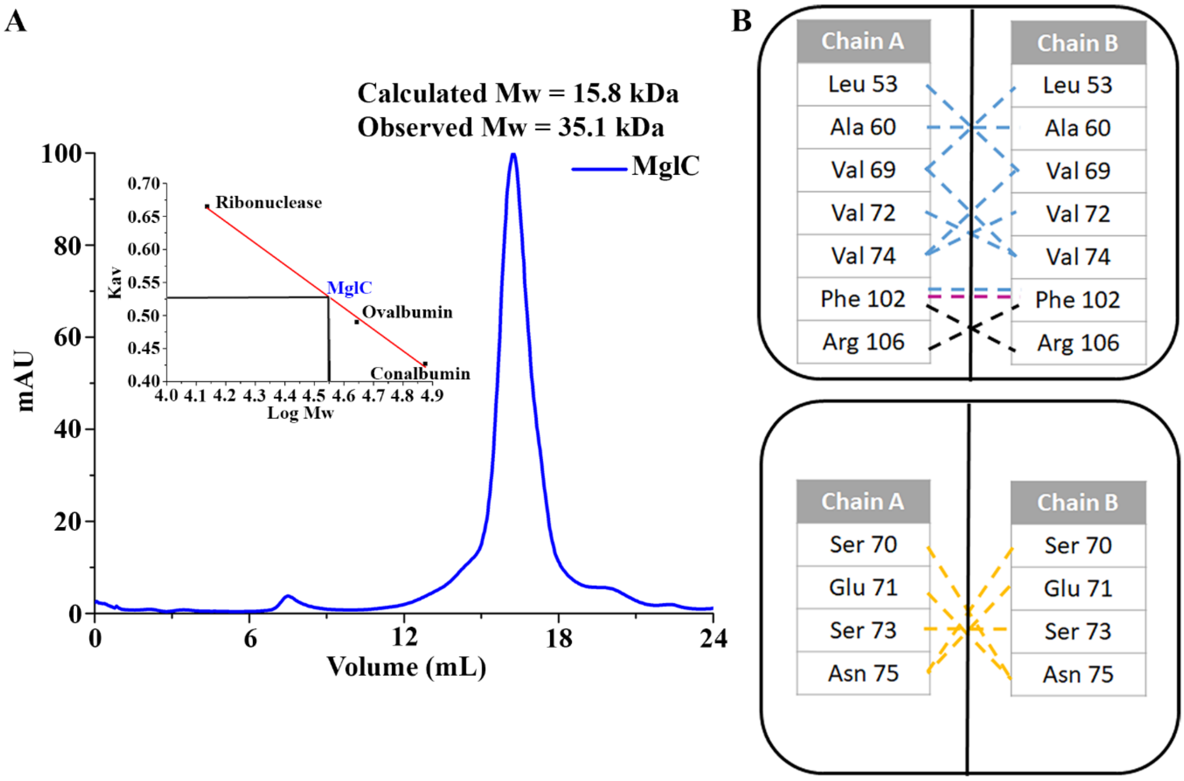


**Supplementary Figure 2: MglC forms dimer (A)** Gel filtration profile of MglC showing that MglC is obligate dimer **(B)** Structural analysis of MglC reveal the residues involved in interaction of dimer. Blue dotted lines represent hydrophobic contacts, yellow dotted lines represent hydrogen bonds, pink dotted lines represent aromatic-aromatic interaction and black dotted lines represent cation-pi interaction (Laskowski et al., 1997).

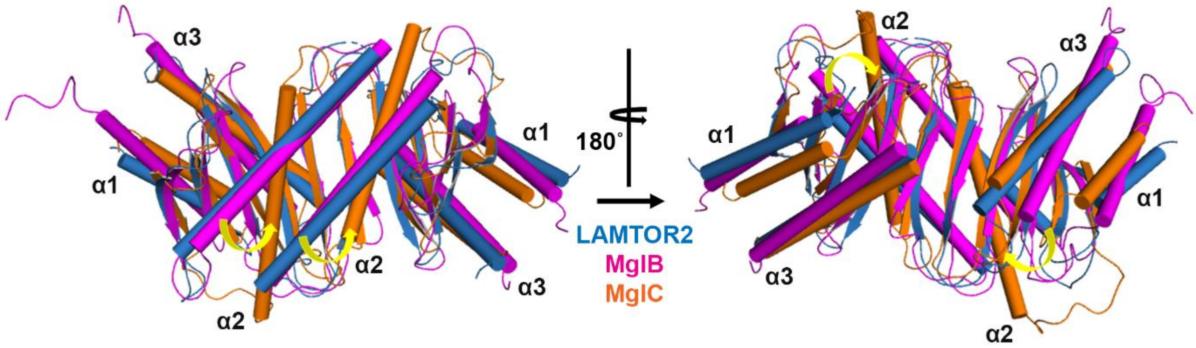


**Supplementary Figure 3: Comparison of MglC with other RLC7 family proteins.** Structural comparison of MglC dimer with the dimers of other RLC7 family proteins showing that in MglC dimer the α1 and α2 are also highly shifted compared to other RLC7 proteins.

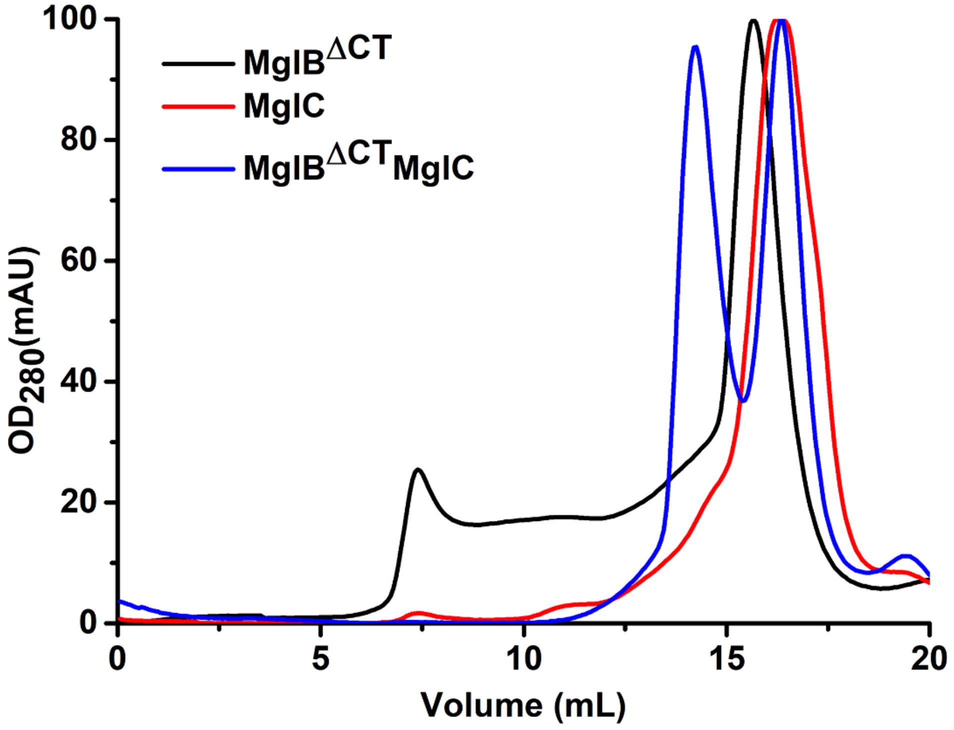


**Supplementary Figure 4: Analytical gel filtration profile suggests that MglB$^{\Delta CT}$ interacts with MglC. (A)** Purified MglB$^{\Delta CT}$ and MglC were mixed and run on the analytical gel filtration column. The gel filtration profile of TEV cleaved MglB$^{\Delta CT}$ (Black), TEV cleaved MglC (Red) and after mixing MglB$^{\Delta CT}$ and MglC (Blue) is shown. Blue line is having two peaks corresponding to MglB$^{\Delta CT}$-MglC complex and MglC. This indicates that MglB$^{\Delta CT}$ also interacts with MglC.

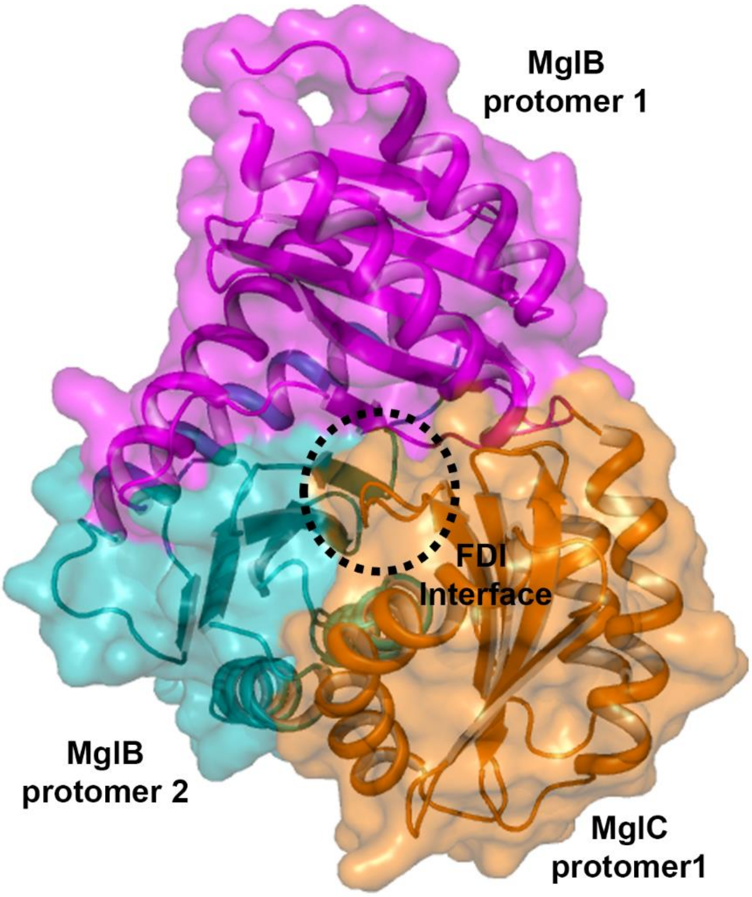


**Supplementary Figure 5: MglC binds MglB with FDI interface.** Model obtained from molecular docking of MglC with MglB shows that FDI interface (Black dotted circle) of MglC is involved in binding MglB.

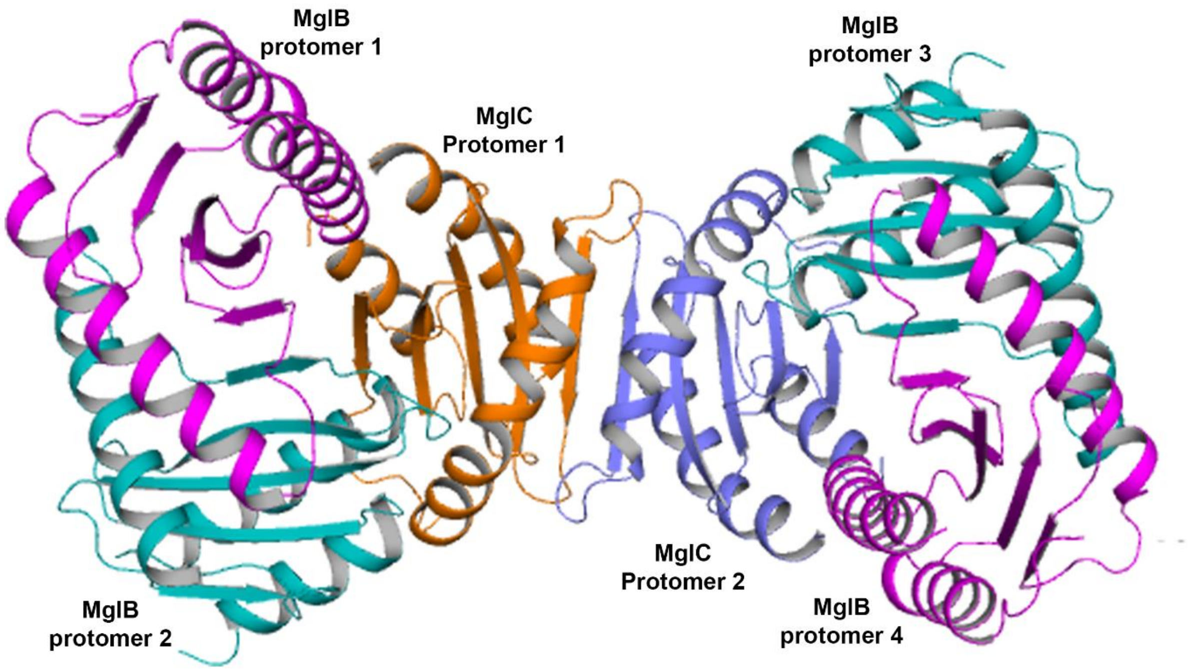


**Supplementary Figure 6: Model for MglBC complex.** MglBC complex model based on experimentally determined binding stoichiometry and docking results.

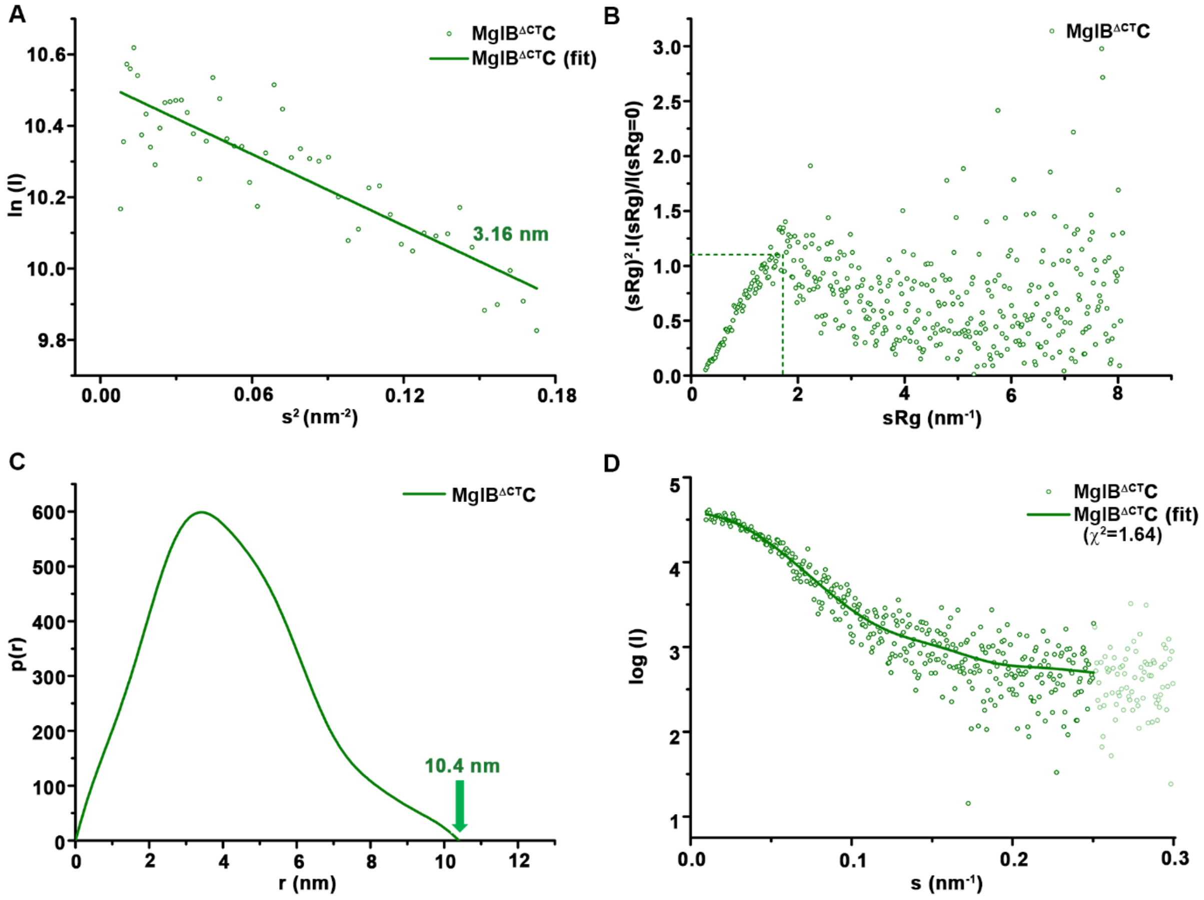

**Supplementary Figure 7: SAXS analysis of MglB$^{\Delta CT}$C. (A)** Guinier analysis of MglB$^{\Delta CT}$C (Rg = 3.16 nm) complex reveals linear fit with no signs of aggregation. **(B)** Dimensionless Kratky plot of MglB$^{\Delta CT}$C complex reveals that the complex is globular as the peak is located at the ideal position as shown by dotted lines. **(C)** Normalized pair distribution function P(r) analysis reveals the $D_{max}$ of 10.4 nm. **(D)** Dummy atom model for MglB$^{\Delta CT}$C complex ($\chi^2$ = 1.64) is prepared using GASBOR (Svergun et al., 2001). The graph represents the intensity profile of MglB$^{\Delta CT}$C complex represent in spheres and the line represents the fitting of dummy atom model generated by GASBOR (Svergun et al., 2001).

**Supplementary Table 1:** Comparative analysis of MglC monomer with other RLC7 family members.

| Protein | Organism | PDB ID : Chain | Z score | r.m.s.d. (Å) | %seq (Sequence identity) | %sse (% of matched secondary structure in target protein) | $N_{align}$ (Number of residues aligned) | Angle of α2 helix compared to MglC α2 helix |
|---|---|---|---|---|---|---|---|---|
| LAMTOR2 -LAMTOR3 (Ragulator complex) | *H. sapiens* | 5y3a : B | 7.8 | 1.88 | 10 | 86 | 94 | 33.39 |
| RLC7 domain | *S. avermitilis* | 3kye : C | 7.8 | 1.77 | 15 | 86 | 94 | 41.12 |
| MglB MglA complex | *M. xanthus* | 6h5b : C | 8 | 1.77 | 9 | 86 | 94 | 41.29 |
| MP1-P14 complex | *H. sapiens*, & *M. musculus* | 1sko : B | 7.4 | 2.01 | 10 | 86 | 94 | 33.64 |
| MglB | *M. xanthus* | 6hjm : D | 7.7 | 1.91 | 9 | 86 | 93 | 37.63 |
| MglB | *T. thermophilus* | 3t1r : A | 3.7 | 3.23 | 13 | 86 | 106 | 35.56 |
| MP1-P14 | N/A | 2zl1 : B | 7.1 | 2.14 | 9 | 86 | 93 | 33.65 |
| LAMTOR2-LAMTOR3 (FLCN-FNIP-Rag-Ragulator complex) | *H. sapiens* | 6ulg : B | 2.9 | 3.3 | 7 | 57 | 101 | 30.79 |
| Human regulator complex | *H. sapiens* | 5yk3 : G | 6.8 | 1.93 | 10 | 86 | 92 | 33.42 |
| Raptor-Rag-Ragulator complex | *H. sapiens* | 6u62 : E | 3.7 | 3.19 | 5 | 86 | 98 | 27.31 |
| Raptor-Rag-Ragulator complex | *H. sapiens* , & *M. musculus* | 5x6v : G | 5.8 | 2.25 | 11 | 71 | 96 | 31.58 |
| Ragulator | *H. sapiens* | 6b9x : B | 3.6 | 3.06 | 5 | 86 | 95 | 35.56 |
| Hepatitis B X-interacting protein | *H. sapiens* | 3msh : A | 6.2 | 2.17 | 11 | 57 | 73 | 34.21 |
| Hepatitis B X-interacting protein | *H. sapiens* | 3ms6 : A | 5.9 | 2.29 | 11 | 71 | 75 | 36.07 |
| MglA in complex with MglB in transition state | *T. thermophilus* | 3t12 : C | 3.9 | 3.19 | 13 | 86 | 106 | 31.45 |

**Supplementary Table 2:** Comparative analysis of MglC dimer with other RLC7 family members. The structural alignments and r.m.s.d. calculations were performed using PyMOL.

| Protein | Organism | PDB ID | r.m.s.d. (Å) | Number of residues aligned |
|---|---|---|---|---|
| LAMTOR2 -LAMTOR3 (Ragulator complex) | *H. sapiens* | 5y3a | 5.11 | 192 |
| RLC7 domain | *S. avermitilis* | 3kye | 7.34 | 136 |
| MglB MglA complex | *M. xanthus* | 6h5b | 5.89 | 176 |
| MP1-P14 complex | *H. sapiens,& M. musculus* | 1sko | 4.54 | 208 |
| MglB | *M. xanthus* | 6hjm | 5.29 | 192 |
| MglB | *T. thermophilus* | 3t1r | 5.57 | 216 |
| MP1-P14 | N/A | 2zl1 | 4.81 | 200 |
| LAMTOR2-LAMTOR3 (FLCN-FNIP-Rag-Ragulator complex) | *H. sapiens* | 6ulg | 5.03 | 208 |
| Human regulator complex | *H. sapiens* | 5yk3 | 5.5 | 200 |
| Raptor-Rag-Ragulator complex | *H. sapiens* | 6u62 | 4.54 | 192 |
| Raptor-Rag-Ragulator complex | *H. sapiens* , & *M. musculus* | 5x6v | 6.38 | 184 |
| Ragulator | *H. sapiens* | 6b9x | 5.07 | 208 |
| Hepatitis B X-interacting protein | *H. sapiens* | 3msh | 5.26 | 88 |
| Hepatitis B X-interacting protein | *H. sapiens* | 3ms6 | 4.28 | 120 |
| MglA in complex with MglB in transition state | *T. thermophilus* | 3t12 | 5.23 | 216 |
